# Water and nutrient cycling regulation functions in forest ecosystems: a comparison between native forests and exotic fast-growing tree species plantations in tropical highlands

**DOI:** 10.1101/2020.04.13.038976

**Authors:** Juan Diego León-Peláez, Juan Camilo Villegas, Jorge Alejandro Amador-Pérez

## Abstract

An improved understanding of ecosystem functions is increasingly needed as ecosystem management moves towards optimizing their capacity to provide services to society. Such a task requires the characterization of ecosystem functions in strategic systems such as tropical mountain forests, which are also subject to pressure due to both global and local environmental changes. In particular, transformation of native forests into pastures or agriculture, has been regarded as the type of transformation with the largest effects in ecosystem regulating and provision functions. However, the effects of other transformations such as those associated with replacement of native with planted forests, have been less studied. To evaluate the effect of forest type on key-ecosystem functions related to water resources, we studied the dynamics of rainfall partitioning and nutrient circulation on a suite of representative forest types in neotropical mountain systems: two plantations of exotic fast-growing species and two types of native forests. Our results illustrate that, when considered in a per-basal unit area, water transmission to the forest floor is significantly higher in both native forests. Similarly, native forests are more effective on circulating nutrients on the ecosystem as they are better adapted to oligotrophic soils such as those occurring in tropical mountains. These results suggest that the replacement of native forests with exotic tree plantations can potentially impact hydrological regulation and the nutrient cycling in these high Andean lands, affecting both directly and indirectly the capacity of ecosystems to produce services to society.

## INTRODUCTION

Tropical mountain systems have a fundamental interest regarding ecosystem services (Briner and others 2013), as a large portion of the population in the tropics lives and depends on these systems for their livelihoods, they share a large proportion of the agricultural productivity, and have been identified as key on regulating ecosystem functions. In particular, tropical mountain systems are essential for maintaining hydrologic regulation functions and nutrient cycling, relevant for water supply for human activities (Locatelli and others 2014, Labrière and others 2015). This is particularly sensitive in the Neotropics, where forest transformation to other land uses is fast, and is still undergoing, with deforestation rates increasing recently (Salazar and others 2018). These areas, although important, are subject to pressure due to both global and local environmental changes (Tropek and others 2014, Balthazar and others 2015). In particular, deforestation—transformation of natural forested ecosystems into pastures or agriculture—has been regarded as a land cover transformation that has profound effects in ecosystem regulating and provision functions (Rounsevell and others 2006, Wasige and others 2013, Suescún and others 2017, García-Leoz and others 2018). In fact, such transformations of vegetation cover, together with climatic variability, may threaten the sustainability of ecosystem functions, as the natural dynamics of different ecosystem processes are strongly affected (Li and others 2009).

Numerous studies have shown the influence of the forest canopies on water and nutrient fluxes to and into soil, modifying the amount and quality of partitioned rainfall (Tan and others 2018). In Andean montane forests, rainfall partitioning (into throughfall, stemflow and interception) and nutrient concentrations in rainfall determine key aspects of nutrient cycling and plant nutrition (Macinnis-Ng and others 2012). Specifically, two considerations regarding the nature and functioning of these forests make them strategic ecosystems of great fragility for the sustainability of ecosystem services: (1) their spatial location in the headwaters with strong anthropic pressure (Ramírez and others 2014), and (2) the extremely nutrient-poor soils on which they are established (Peláez-Silva and others 2018). Therefore, needed are assessments of the potential effects in both the water balance and water chemistry to better assess the effects of native forest transformations into other vegetation cover types (Tan and others 2018).

Among the potential transformations of native forests, perhaps the least studies are the effects of other transformations such as those associated with replacement of native with planted forests (Mooney and others 2009, Suescún and others 2018). In particular, the use of fast-growing, exotic tree species for the establishment of those plantations, has been the subject of heated debate for a long time (Peláez-Silva and others 2018). More specifically, in countries with high biodiversity, there are potentially negative consequences associated with such plantations, including alteration of successional processes (D’Antonio and Meyerson 2002), reduction of biodiversity (Pawson and others 2013), effects on water availability (Bonnesoeur and others 2019) and nutrient cycling processes (Ramírez and others 2014) in watersheds. These effects imply the potential alteration of the supply and impoverishment of water and soil quality, related to ecosystem regulation functions (e.g. water and nutrient cycling functions) fundamental for the provision of key ecosystem services.

These aspects are of special relevance in countries with recognized forestry potential, as is the case of Colombia, where the lands officially identified for commercial tree planting (Fajardo-Mejía and others 2016), comprise about 24 million hectares (ca. 21% of the country). These include Andean highlands and the use of fast-growing exotic tree species, whose effects are still poorly known in terms of the provision of water-related ecosystem services. Consequently, land managers and decision makers are challenged when considering the selection of species for multiple purposes, and when evaluating the ecological and environmental effects of those decisions (Daily and others 2009). Therefore, an improved understanding ecosystem functions and their variations between forest types, is increasingly needed as management of forest ecosystem services moves toward maintenance-based optimization. Such a task requires the characterization and appropriate differentiation of ecosystem functions in strategic systems such as tropical mountain areas.

As an approach to evaluate the effect of forest type on key-ecosystem functions related to water resources (quantity and quality), we analyzed the dynamics of rainfall partitioning and nutrient cycling on a suite of representative forest types in Andean highlands: (1) two types of native forests (a mixed-composition secondary growth and an oak -*Quercus humboldtii*-dominated system) and (2) two types of planted forests established with fast-growing exotic species (*Pinus patula* and *Cupressus lusitanica*). To do this, we monitored incoming precipitation, throughfall, and stemflow and determined nutrient flows in all of them, using a space for time substitution approach (García-Leoz and others 2018), in which the four forest types were monitored at the same time. We highlight the potential effects of forest type on the sustainability of key ecosystem services related to water in tropical mountain areas.

## MATERIALS AND METHODS

### Study Site

Field research was conducted over the course of one year in the Piedras Blancas watershed, which is located in the Nare Protected Forest Reserve in the central Andes of Colombia (Peláez-Silva and others 2019). Mean annual precipitation is 1936 mm, mean annual air temperature ranges between 14.9 and 16.0 °C, and the mean annual relative humidity is 83%. The area is located within the lower montane humid forest, according to the Holdridge life zone classification. The landscape is dominated by low- to mid-slope hills covered by volcanic ash. Soils are classified as Fulvudands and Hapludands. These soils have good physical properties, generally acid and with high contents of organic matter, generally very low in major elements and available nitrogen, and with phosphorous fixating properties.

### Study plots

To describe the differential effects of forest type in the dynamics of canopy water fluxes, we experimentally evaluated the dynamics of rainfall partitioning and nutrient circulation on a suite of representative forest types in Andean highlands (Table 1): (1) two types of native forests (a mixed-composition secondary growth and an oak -*Quercus humboldtii*-dominated system); and (2) two types of planted forests established with fast-growing exotic species (*Pinus patula* and *Cupressus lusitanica*). In order to compare the hydrological and biogeochemical processes between these types of forests, a study area as uniform as possible was selected (ca. 40 ha, 2460 masl, 6 ° 15’42‘‘N, 75 ° 30’30 ‘‘W) where the four ecosystems were represented, so that the conditions of the physical environment were very similar for them (ie topography, soil, climate). Tree plantations were established 40 years prior to the study without the use of silvicultural practices; nevertheless they have been subjected to furtive individual tree withdrawals. Before these plantations, the soils were severely deforested and degraded as a result of forest conversion to grassland for cattle. The oak forest represents the predominant Colombian montane forest type (Peláez-Silva and others 2018), which is dominated by *Quercus humboldtii*. The other native forest is a secondary forest where predominant species include *Clethra fagifolia, Clusia discolor, C. multiflora, Croton magdalenensis, Oreopanax floribundum, Tibouchina lepidota, Vismia baccifera, V. ferruginea, Weinmannia balbisiana, Befaria aestuans, Palicourea angustifolia*.

**Table 1.** Structural characteristics for native forests and coniferous plantations at the Piedras Blancas watershed.

| Forest | Age<br>(years) | N<br>(ind ha<sup>-1</sup>) | Dq<br>(cm) | H<br>(m) | G<br>(m<sup>2</sup> ha<sup>-1</sup>) |
| --- | --- | --- | --- | --- | --- |
| Cypress plantation | 40 | 260 | 22,9 | 22,7 | 21,4 |
| Pine plantation | 40 | 310 | 36,7 | 24,6 | 65,7 |
| Secondary forest | 20 | 540 | 12,5 | 12,7 | 9,4 |
| Oak forest | >60 | 347 | 20,2 | 17,0 | 13,2 |

N: stand density for trees with dbh >10 cm, Dq: mean tree diameter squared, H: mean canopy height, G: stand basal area for all trees with dbh >10 cm

### Field instrumentation and monitoring

Three cylindrical 6-inches-diameter rain gauges were located in the open field for incoming precipitation (Pi) measurement. On each forest type, additional ten cylindrical rain gauges (like those used for Pi) were installed for throughfall collection (Th), which corresponds to the amount of rain that reaches the soil after passing through forest canopies. Since the rain gauges occupy a specific spatial position, and that the distance between them constitutes the factor that can determine the condition of independence or dependence given their association to different microsite conditions, their distribution, as sample units, was made based on a randomized restricted scheme. For this, it was controlled that the distance between any pair of units was at least 15 m, which, although not guaranteeing the obtaining of independent readings, does tend to do so (Marriot and others 1997). For stemflow (Sf) measurement, which corresponds to the amount of water that drains around tree trunks, we installed ten stem collars for each type of forest. The location of the stem collars was made based on a stratified sampling, taking the tree diameter distribution as a stratification criterion, establishing the number of units proportional to the size of the stratum. Each stem collar was arranged helically in each tree, guaranteeing the total coverage of its diameter. Water collected in all the devices was conducted to plastic jars. Volumetric measurements were taken weekly for one year. With a monthly frequency, combined samples of all devices on each hydrological flow in all forest types were taken for chemical analysis in the laboratory.

### Chemical analysis

Monthly combined samples of Pi, Th, and Sf were filtered and analysed for pH, calcium, magnesium, potassium, phosphate, and nitrogen concentrations (mg L^-1^). The determination of calcium, magnesium, and potassium was performed by direct reading through atomic absorption spectrophotometry. The determination of total N was made by means of photometric analysis (NANOCOLOR®), prior decomposition in digester block (oxidative decomposition) and reading at 365/385 nm. The determination of P (PO_4_-P) was carried out following the ascorbic acid method, being read in a visible ultraviolet spectrophotometer (Bausch and Lomb Milton Roy 601) at 660 nm. The analysis were carried out in the Biogeochemistry Laboratory of the National University of Colombia, Medellín.

### Data processing and statistical analysis

Weekly volumes of Pi and Th were converted to water depth (mm), dividing them by the collection area of the rain gauges. To extrapolate Sf measurements from tree-to-forest, we considered the volume of water collected and the basal area of the trees that had collars installed, with respect to the total basal area of the stand (as described in García-Leoz and others 2018). The Net Precipitation (Np) was calculated as Np = Th + Sf. Interception was calculated as I = Pi-Np. The Th and Sf water depths were compared by ANOVAS, after verification of the normality of the data (Shapiro-Wilk test). The differences between means were determined by the Tukey test (*P=*0.05). When the variables were not normally distributed, the Kruskal-Wallis test (*P*=0.05) was used.

To determine how nutrients circulate within the canopies, we used the concentration ratio (Cr), which was calculated as the ratio between the mean volume-weighted nutrient concentrations in the Th and the mean volume-weighted nutrient concentrations in the Pi (Parker 1983). Nutrient flows (kg ha^-1^ year^-1^), were obtained from the product between the mean volume-weighted nutrient concentrations (mg L^-1^) and the volumes of each flow. The net contributions of nutrients, which include processes occurring in the canopy and in the path of the water until reaching the forest floor (washing of the material deposited in the canopy or dry deposition, leaching of ions from the inside of plant organs, absorption of ions from precipitation into plant structures), were determined through the Net Deposition (Nd). This was calculated as the difference between the amount of the nutrient in the net precipitation and the amount of that nutrient in the rainwater (Nd=Np-Pi). In addition, to indicate the relative gain or loss of mass of nutrients of the incoming precipitation after passing through the canopy, the Deposition Index proposed by Parker (1983) (Di=Np/Pi) was calculated.

Given that the forests were structurally different, we normalized hydrological and hydrochemical flows, by the degree of occupation of the forest stand using the basal area (G, m^2^ ha^-1^) as a way to standarize Np values (mm=L m^-2^). Thus, the Np/G ratio was obtained for each forest, expressed as m^3^ water/m^2^ tree for hydrological flows, and as kg nutrient/m^2^ tree for hydrochemical flows. Aditionally, we considered the Np/G ratio values obtained for the flows of each nutrient in each of the native forests, as reference values for comparison with those corresponding to the coniferous plantations. These relative values (Np/G)_rel_ were calculated for each nutrient from the division between the Np/G value corresponding to the coniferous plantation, and the Np/G value corresponding to the native forest, expressing the result as percentage.

## RESULTS

### Canopy water flows

The annual Pi accounted for 2074 mm, with maximum and minimum absolute values in the first (July-August 2008: 292 mm) and tenth month (May-June 2009: 78 mm), respectively (Figure 1). Based on the differences between Np and Pi (Figure 1), extra water income was determined in the forests studied (cases in which net precipitation was higher than incoming precipitation). The number of weeks in which Np>Pi followed the pattern: oak forest (35)> pine plantation (15)> secondary forest (11)> cypress plantation (7). Following this order, excess precipitation during the entire record was: oak forest (343) mm)> pine plantation (113) mm> secondary forest (77 mm)> cypress plantation (28 mm). These values represented 16.5% (oak forest), 5.4% (pine plantation), 3.7% (secondary forest) and 1.3% (cypress plantation) of the total incoming precipitation.

**Figure 1.**
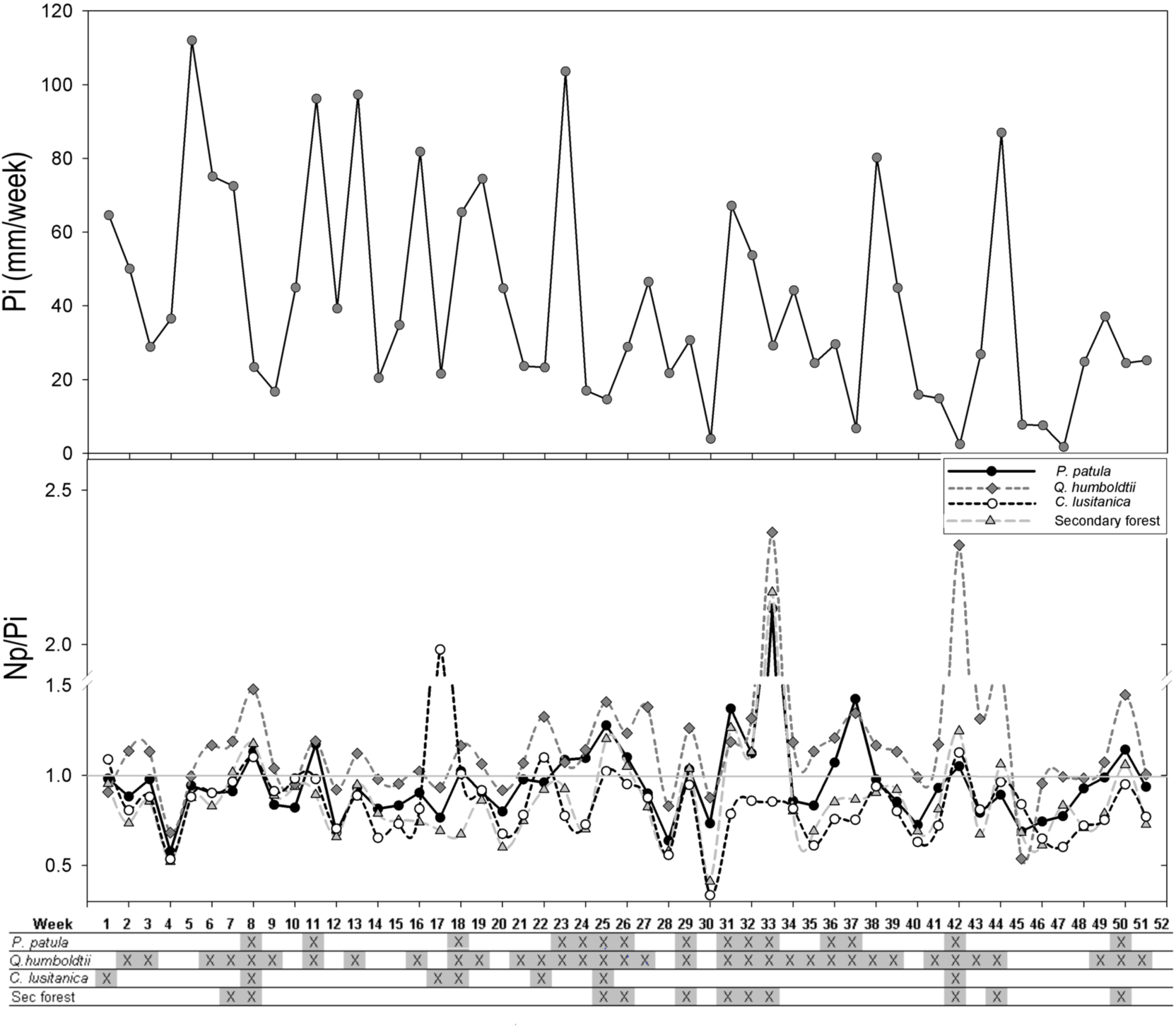
Weekly values of incoming precipitation (Pi, mm) and net precipitation/incoming precipitation ratio (Np/Pi) in the four forests during the study period. Letter X in the lower part of the figure indicate for each week and forest, the occurrence of values Np> Pi, indicative of potential water addition by fog interception.

The highest Interception (I) was found in the secondary forest and in the cypress plantation (ca.15% of the Pi), without significant differences between them (*P*=0.3518). The Th in the four forests represented more than 85% of the Pi (Figure 2), with no significant differences between their weekly values (*P*=0.9013) and showing a close linear correlation with the Pi (r Pearson=0.95-0.98, *P*=0.05). Sf had a very low participation in the overall water budget, being highest in the pine plantation (25 mm, 1.2% of the Pp) and lowest in the oak forest (2.9 mm, 0.2% of the Pp). Significant differences were found (*P*<0.05) in the Sf between the oak forest and the other forest types, which presented higher Sf.

**Figure 2.**
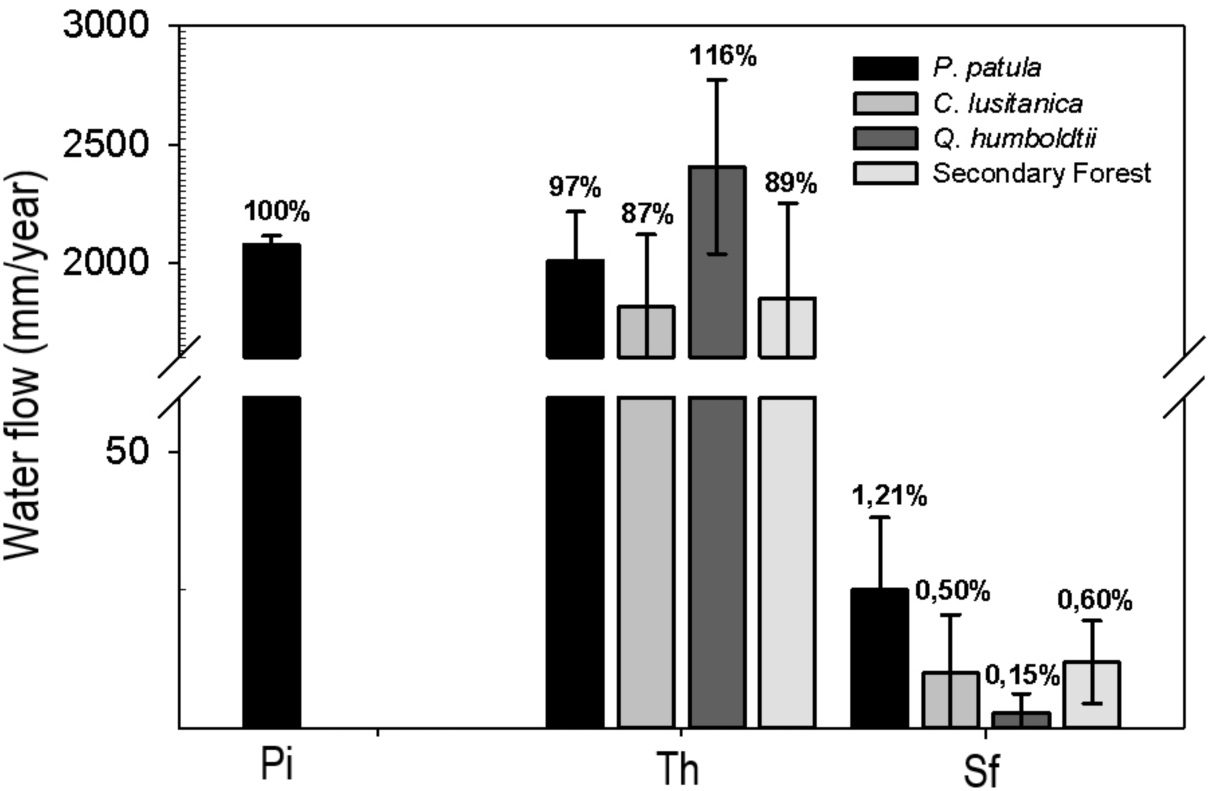
Annual incoming precipitation (Pi) and its partitioning into throughfall (Th) and stemflow (Sf) in the four forests during the study period. Values expressed as water depth (mm) and as percentage (%) with respect to the total Pi.

### Canopy water flow chemistry

The mean pH of the Pi was 5.6, characterized by a low temporal variation (CV=16.1%), showing changes when coming into contact with the canopy. Thus, the mean pH value for Th was higher than that of Pi in the cypress plantation (6.1) and in the oak forest (5.7), obtaining its lowest value in the pine plantation (5.2). pH values of Sf with respect to precipitation were higher in all the forests (average pH=6.2), except for pine plantation where the lowest value was found (pH=4.5).

Nutrient concentrations in Pi followed the decreasing sequence: N>K>Ca>Mg=P (Figure 3). For Th, in general, the highest nutrient concentrations were found in the secondary forest, and the lowest in the pine plantation. The highest nutrient concentrations in this flow were found for N (1.16-1.83 mg N L^-1^) and the lowest for P (0.05-0.10 mg P L^-1^). There were no significant differences among forests for the same nutrient in the same hydrologic flux (Th and Sf; Kruskal-Wallis, *P*> 0.05). The majority of nutrients showed increases in their concentrations when moving from Pi to Th, except for P, with values reducing in both conifer plantations (Cr<1.0, Figure 3). With the exception of N, nutrient concentrations increased from Th to Sf. In general, the highest concentrations in the Sf occurred in the cypress plantation and the lowest in the pine plantation, with K being the nutrient with the highest concentrations (Figure 3).

**Figure 3.**
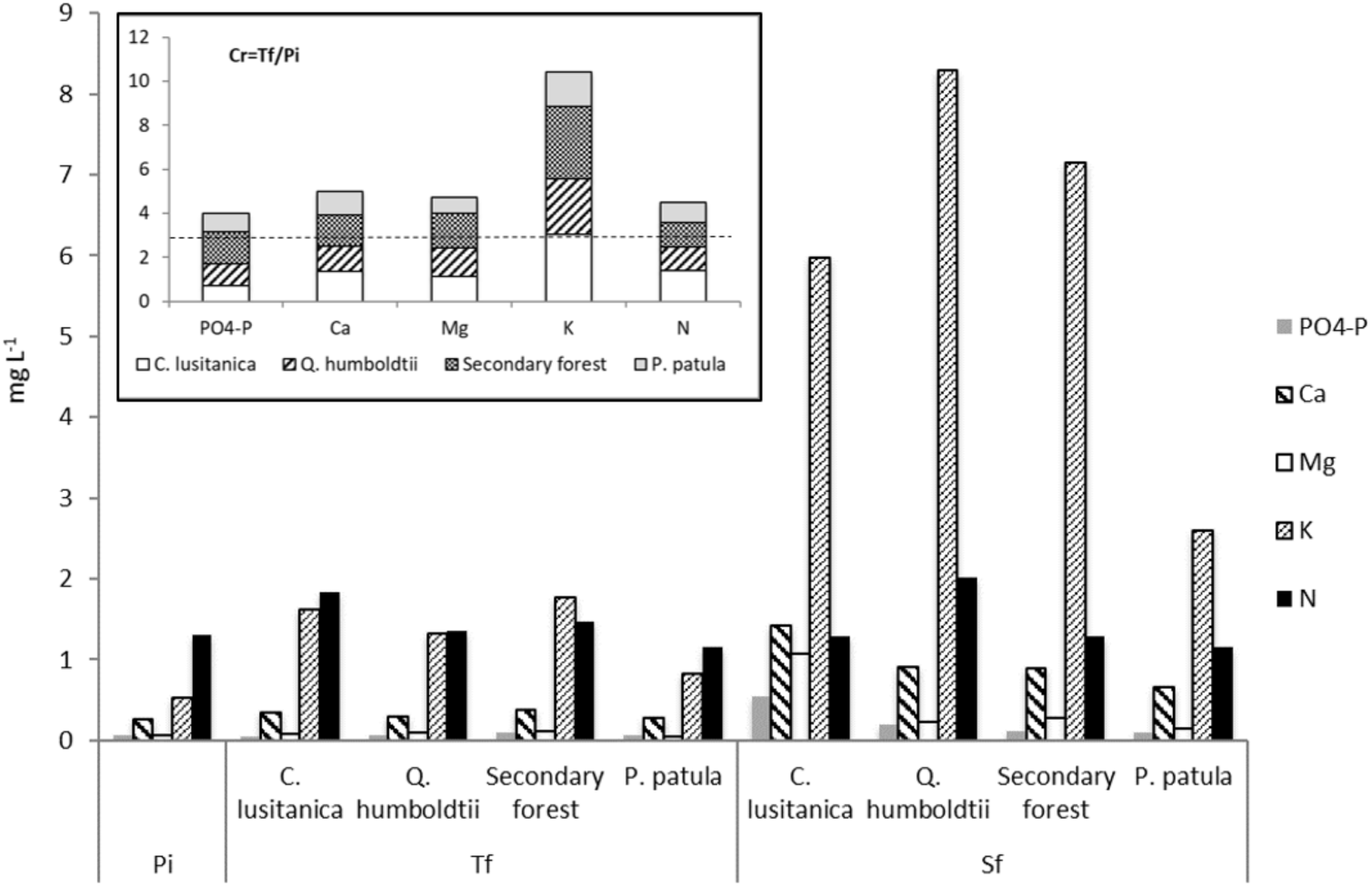
Mean volume-weighted concentration in the incoming precipitation (Pi), throughfall (Th) and stemflow (St) flows, and values of the concentration ratio Cr (Inner graph) in the four forests, where the horizontal line represents the threshold for enrichment or impoverishment from incoming precipitation, corresponding to a concentration ratio value of 1.0.

### Nutrient deposition and nutrient canopy exchange

The highest total nutrient input through precipitation (Pi) occurred with N, followed by K and Ca (Table 2). The highest flow of nutrients via Th occurred in the oak forest, however no significant differences were found for any nutrient. In Sf, the highest values were found in the pine plantation, with the exception of Mg, with its highest value occurring in the cypress plantation. N was the highest nutrient in the Np of all the forests, followed closely by K. Although no significant differences were detected among forests in the nutrient flows via Np (Kruskal-Wallis, *P*> 0.05), the highest values were found in the oak forest, consistent with previously noted trends. For the set of nutrients studied here (Ca+Mg+K+P+N), the annual amounts contributed by the Np followed the sequence: oak forest (71.5 kg ha^-1^)> cypress plantation (55.6 kg ha^-1^)> secondary forest (53.3 kg ha^-1^)> pine plantation (39.4 kg ha^-1^). Net Deposition (Nd, Figure 4), showed positive and negative values with no particular specific trend. Notably, Nd was positive for all nutrients only in the oak forest, highlighting nutrient enrichment in water as it passes through the canopy. Consistent positive values of Nd occurred in all four forest types only for K. In the oak forest, these gains in K represented, in relative terms (Di values, Figure 4), increases three times higher with respect to the quantities of the nutrient contributed by the Pi.

**Table 2.** Inputs and annual nutrient flows via incoming precipitation (Pi), throughfall (Th), stemflow (Sf) and net precipitation (Np) in the four forests.

| Flow | Forest | PO<sub>4</sub>-P | Ca | Mg | K | N |
| --- | --- | --- | --- | --- | --- | --- |
| Pi | - | 1,35 | 5,20 | 1,33 | 9,12 | 21,61 |
| Th | Pine plantation | 1,2 a | 4,57 a | 0,83 a | 11,69 a | 20,31 a |
|  | Oak forest | 1,54 a | 6,13 a | 1,63 a | 28,98 a | 33,11 a |
|  | Cypress plantation | 0,87 a | 4,60 a | 1,07 a | 22,07 a | 26,62 a |
|  | Secondary forest | 1,86 a | 3,48 a | 1,10 a | 21,41 a | 25,35 a |
| Sf | Pine plantation | 15,98 a | 89,98 a | 20,65 a | 415,95 a | 218,64 a |
|  | Oak forest | 2,22 b | 11,24 b | 2,93 b | 111,17 b | 20,68 b |
|  | Cypress plantation | 12,19 a | 41,49 a | 24,80 a | 165,53 c | 38,22 a |
|  | Secondary forest | 1,30 b | 9,22 b | 3,23 b | 77,86 b | 13,31 b |
| Np | Pine plantation | 1,22 a | 4,66 a | 0,85 a | 12,11 a | 20,53 a |
|  | Oak forest | 1,54 a | 6,14 a | 1,63 a | 29,09 a | 33,13 a |
|  | Cypress plantation | 0,88 a | 4,64 a | 1,09 a | 22,24 a | 26,66 a |
|  | Secondary forest | 1,86 a | 3,49 a | 1,10 a | 21,49 a | 25,36 a |

Values for Pi, Th, and Np expressed in kg ha<sup>-1</sup> year<sup>-1</sup>, and for Sf in g ha<sup>-1</sup> year<sup>-1</sup>. Different letters for a nutrient within the same flow (Th, Sf, Np) denote significant differences among forests (Kruskal-Wallis test, P<0.05).

**Figure 4.**
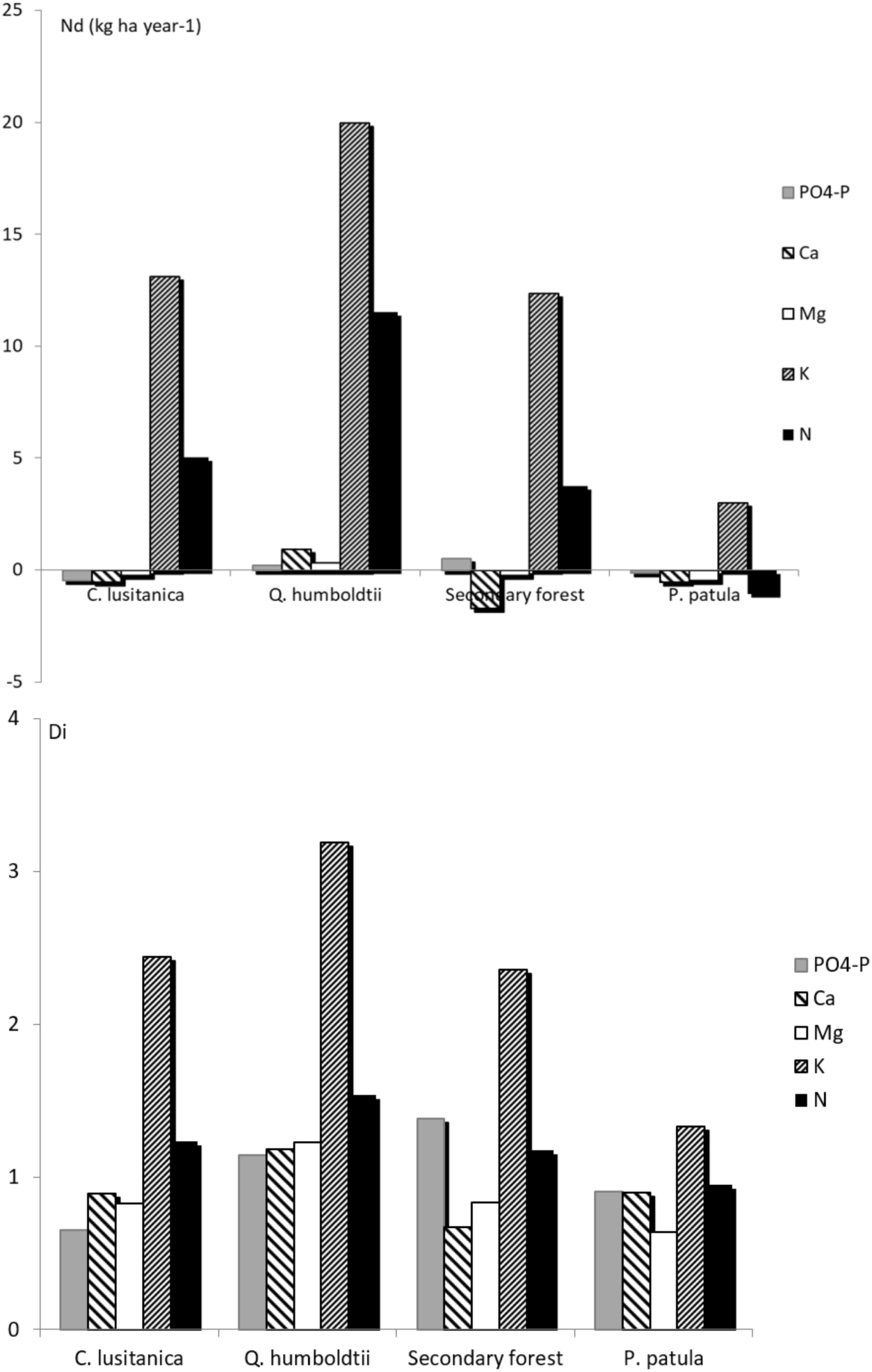
Annual values of net deposition (a) (Nd: kg ha^-1^ year^-1^) and deposition index (Di) (b) of nutrients in the four forests. The net deposition index (Nd) was calculated for each nutrient as Nd=net precipitation (Np)-incoming precipitation (Pi). The deposition index (Di) was obtained for each nutrient as Di=net precipitation (Np)/incoming precipitation (Pi).

### Hydrological and hydrochemical regulation

Given that our forests (both the natural and the planted forests) exhibited significant structural differences, and that forest structure influences hydrological and hydrochemical function, we propose a regulation metric, based on forest occupation, as explained in the previous section. Even as the oak forest showed a high capacity for capturing extra water input (a high Np value), when comparing the Np/G ratio values for the hydrological flows of the four forests studied here, the secondary forest exhibited a higher Np/G value (1980 m^3^ water/m^2^ tree), yet similar to that of the oak forest (1820 m^3^ water/m^2^ tree), which has a higher basal area. Beyond the slight differences between these two natural ecosystems in the Np/G ratio, they are notably higher than Np/G values in the planted forests (Pine=310, Cypress=850), indicating a higher regulatory capacity per occupation unit. Similarly, native exhibited higher capacity to regulate hydrochemical flows based on the annual values of the Np/G ratio. Thus, the mean Np/G values found in both the native forests and the coniferous plantations for the different nutrients were, respectively (kg of nutrient/m^2^ tree): P (0.16 and 0.03); N (2.60 and 0.78); Ca (0.42 and 0.14); Mg (0.12 and 0.03); and K (2.1 and 0.61).

Relative Np/G values – (Np/G)_rel_ – obtained from comparing nutrient flows between native forests and coniferous plantations (Figure 5), show that in most cases, nutrient flows in coniferous plantations were below 50% of those corresponding to native forests. The lowest values were found in the *P. patula* plantation, in which, nutrient flows where always lower than 20% of the values found in native forests. Although in the *C. lusitanica* plantation nutrient flows were higher (compared to *P. patula*), only in two nutrients flows were equal or greater than 50% (Ca when compared with the secondary forest and N when compared with the oak forest; underlined values in Figure 5).

**Figure 5.**
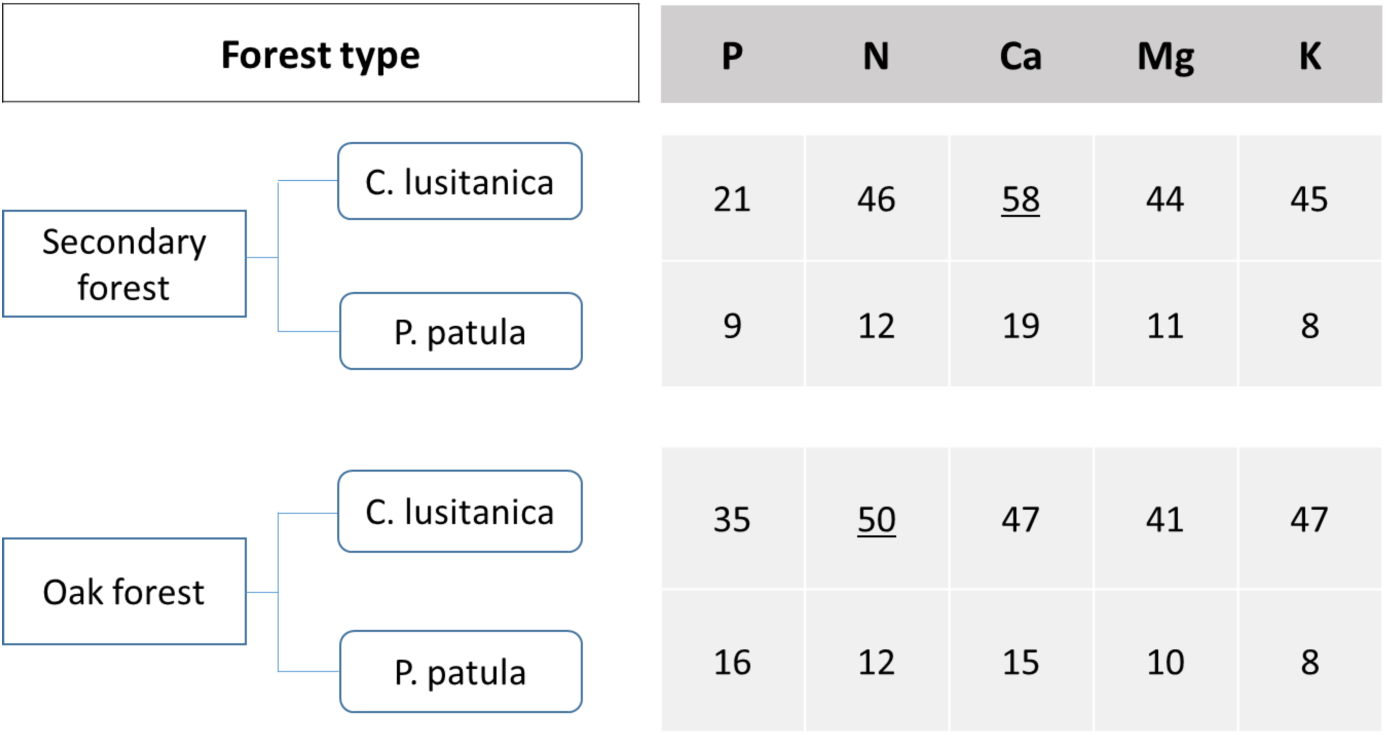
Comparison of the (Np/G)_rel_ values (%) obtained for each nutrient between native forests and coniferous plantations. Each value represents the percentage reached of the Np/G ratio for a specific nutrient, compared to each specific type of forest. Values obtained in plantations equal to or greater than 50% of the values obtained in native forests are underlined.

## DISCUSSION

### Canopy water fluxes

Our results suggest that natural forests could be more effective in transmitting water fluxes from the canopy into the forest floor. This was particularly observed in the oak forest, with consistent higher values of net precipitation than incoming precipitation (although occurring less frequently in all forests), suggesting the potential occurrence of horizontal precipitation (or fog deposition), that led to an overall increase in hydrological input of 16%, compared to incoming precipitation (Figure 2). As a consequence, Th values obtained for these four forests were located at the upper limit of the reports for other tropical montane forests, while Sf did not represent a significant contribution to the overall forest water budget (as highlighted in previous studies, Macinnis-Ng and others 2012). However, in our study, the Sf flux differed in the oak forest from all other forest types. Overall, our results highlight the role of natural forests, particularly oak-dominated forests (generally associated with mature and well-preserved stages in tropical montane forests of south America). More specifically, these forests favor the movement of water through the canopy, therefore suggesting a potential improvement in ecosystem functions that result in key-ecosystem services.

### Chemical alteration of incoming precipitation

Changes in the chemical composition of water that reaches the forest floor (as throughfall-Th) are generally the result of the interaction of incoming precipitation (Pi) with the canopy (Scheer 2011). Nutrients from precipitation that reach the canopy change as the result of two processes that include the washing of dry deposition accumulated on the canopy, and canopy exchange through nutrient leaching and direct uptake from the canopy (Tan and others 2018). When comparing nutrient concentrations of the Pi with those of the Th, using the concentration ratio metric (Cr in Figure 3), a general pattern of enrichment was found for K and N when passing the canopy (Cr>1), similar to patterns found in long-term studies (Heartsill-Scalley and others 2007). In contrast to the coniferous plantations, where P was found to be impoverished, concentration was maintained in the oak forest and enriched in the secondary forest. The maximum nutrient enrichment was found for K, as a result of its high mobility and concentration in foliar tissues (Sardans and Peñuelas 2015). In general, both the concentrations and the flows of Ca, Mg and K in the Th and Sf were located in the lower limits of the reports for other tropical montane forests (Cavelier and others 1997, Hölscher and others 2003, Suescún and others 2018), while for N we found the opposite. Although foliar leaching is important in oligotrophic systems, by means of foliar capture, nutrients of low soil availability are removed as P (Jordan and others 1980), as it was found in pine and cypress plantations (Cr<1).

Canopy exchange and washing of the dry deposition represent important nutrient transfer processes within forest ecosystems (Tan and others 2018). Consequently, Np (Th + Sf) has been recognized as an important flow path in the internal dynamics of nutrients worldwide (Macinnis-Ng and others 2012). These results are particularly relevant since such annual returns of key nutrients as N and P were even higher than those from the decomposition of leaf litter in oak forests in the region (León and others 2011). More generally, the higher total amount of nutrients, via Np, that reached the forest floor in the oak forest, represent a fundamental biogeochemical aspect of ecosystem function that is lost when native forests are replaced by planted ecosystems.

In absolute terms, net nutrient inputs (Nd) showed both gains (joint effects of foliar leaching and washing of dry deposition accumulated on the canopy) and net decreases (foliar uptake in the canopy). Positive Nd values found for all the nutrients in the oak forest, indicated an enrichment of the Pi after its contact with the canopy. For N and P, which are recognized as limiting nutrients in these tropical mountains (León and Osorio 2014), the contributions found in the native forests are highlighted, particularly those of N in the oak forest (Nd = 11.5 kg N ha^-1^ year^-1^, Di = 3.2) and those of P in the secondary forest (Nd = 0.5 kg P ha^-1^ year^-1^, Di = 1.4). Overall, greater P limitations appear to occur in coniferous plantations than in the native forests studied. Although these limitations likely result from its low availability in the soil due to its strong fixation in these andisols (Peláez-Silva and others 2018), Nd <1 values found in the plantations (unlike the native forests) suggest its absorption in the canopy. The Nd negative values and Di <1 also obtained for Ca and Mg in both plantations, reflected their possible retention and/or absorption in the canopy in response to the strong soil acidity. Consequently, our results, again, suggest that native forests could be more effective than coniferous plantations on nutrient cycling at ecosystem level and better adapted to oligotrophic soils such as those occurring in tropical mountains.

### Hydro-chemical regulation

Both types of native forests showed greater efficiency in hydrological and hydrochemical regulation in these high Andean lands than what was found in both coniferous plantations, as they not only captured extra water input, but also enriched water with nutrients from the canopy (Dezzeo and Chacón 2006, Liu and others 2016, Tan and others 2018). In terms of nutrient mass, and considering our proposed “per unit occupation” metric, both native forests were very effective on transferring nutrients to the soil via net precipitation. This suggests an intense nutrient cycling in these forests, which is associated with improved ecosystem functioning and potentially on productivity, given the scarcity of nutrients characteristic of these soils. Specifically, these nutrient flows reflect ion exchange processes of valuable ecosystem significance for nutrition and forest health (Heartsill-Scalley and others 2007), as it influences the activity of soil microorganisms, and the replenishment of soil nutrients.

Overall our results suggest that the replacement of native forests with exotic tree plantations could impact hydrological regulation and the nutrient cycling in these high Andean lands, affecting both directly and indirectly the capacity of ecosystems to produce services to society. More specifically, the native forests studied here were highly effective on precipitation redistribution and nutrient cycling, showing their adaptation to oligotrophic soils such as those occurring in tropical mountains, which allow them to effectively function and produce ecosystem services. Land managers and decision makers are often challenged when evaluating the effects of native forest replacement. Our results show that not all forest types are equally effective on maintaining key-ecosystem services such as those related to water and nutrient cycling functions of ecosystems. This is a fundamental management aspect in strategic systems such as tropical mountains, on which human society depend for the supply of drinking water and the development of productive activities.

## ACKNOWLEDGMENTS

National University of Colombia, engineers, students, laboratory and field assistants. The authors thank J. Herrera for technical support. The authors do not have any conflict of interests to declare.

## REFERENCES

Balthazar V, Vanacker V, Molina A, Lambin EF (2015) Impacts of forest cover change on ecosystem services in high Andean mountains. Ecol Indic 48: 63–75.

Bonnesoeur V, Locatelli B, Guariguata MR, Ochoa-Tocachi BF, Vanacker V, Mao Z, Stokes A, Mathez-Stiefel S (2019) Impacts of forests and forestation on hydrological services in the Andes: A systematic review. Forest Ecol Manag 433: 569–584.

Briner S, Huber R, Bebi P, Elkin C, Schmatz DR, Grêt-Regamey A (2013) Trade-offs between ecosystem services in a mountain region. Ecol Soc 18: 35.

Cavelier J, Jaramillo M, Solis D, de León D (1997) Water balance and nutrient inputs in bulk precipitation in tropical montane cloud forest in Panama. J Hydrol 193: 83–96.

D’Antonio C, Meyerson LA (2002) Exotic plant species as problems and solutions in ecological restoration: A synthesis. Restor Ecol 10: 703–713.

Daily GC, Polasky S, Goldstein J, Kareiva PM, Mooney HA, Pejchar L, Ricketts TH, Salzman J, Shallenberger R (2009) Ecosystem services in decision making: time to deliver. Front Ecol Environ 7: 21–28.

Dezzeo N, Chacón N (2006) Nutrient fluxes in incident rainfall, throughfall, and stemflow in adjacent primary and secondary forests of the Gran Sabana, southern Venezuela. Forest Ecol Manag 234: 218–226.

Fajardo-Mejía MA, Morales-Osorio JG, Correa-Londoño GA, León-Peláez JD (2016) Effect of plant extracts and growth substrates on controlling damping-off in Pinus tecunumanii seedlings. Cerne 22: 317–324.

García-Leoz V, Villegas JC, Suescún D, Flórez CP, Merino-Martín L, Betancur T, León JD (2018) Land cover effects on water balance partitioning in the Colombian Andes: improved water availability in early stages of natural vegetation recovery. Reg Environ Change 18: 1117–1129.

Heartsill-Scalley T, Scatena FN, Estrada C, McDowell WH, Lugo AE (2007) Disturbance and long-term patterns of rainfall and throughfall nutrient fluxes in a subtropical wet forest in Puerto Rico. J Hydrol 333: 472–485.

Hölscher D, Köhler L, Leuschner C, Kapelle M (2003) Nutrient fluxes in stemflow and throughfall in three successional stages of an upper montane rain forest in Costa Rica. J Trop Ecol 19: 557–565.

Jordan C, Golley F, Hall J (1980) Nutrient scavenging of rainfall by the canopy of an Amazonian rain forest. Biotropica 12: 61–66.

Labrière N, Locatelli B, Laumonier Y, Freycon V, Bernoux M (2015) Soil erosion in the humid tropics: A systematic quantitative review. Agr Ecosyst Environ 203: 127–139.

León JD, González MI, Gallardo JF (2011) Ciclos biogeoquímicos en bosques naturales y plantaciones de coníferas en ecosistemas de alta montaña de Colombia. Rev Biol Trop 59: 1883–1894.

León JD, Osorio NW (2014) Role of litter turnover in soil quality in tropical degraded lands of Colombia. Sci World J 2014: 1–11.

Li Z, Liu WZ, Zhang XC, Zheng FL (2009) Impacts of land use change and climate variability on hydrology in an agricultural catchment on the Loess Plateau of China. J Hydrol 377: 35–42.

Liu J, Zhu L, Wang H, Yang Y, Liu J, Qiu D, Ma W, Zhang Z, Liu J (2016) Dry deposition of particulate matter at an urban forest, wetland and lake surface in Beijing. Atmos Environ 125: 178–187.

Locatelli B, Imbach P, Wunder S (2014) Synergies and trade-offs between ecosystem services in Costa Rica. Environ Conserv 41: 27–36.

Macinnis-Ng CMO, Flores EE, Müller H, Schwendenmann L (2012) Rainfall partitioning into throughfall and stemflow and associated nutrient fluxes: land use impacts in a lower montane tropical region of Panama. Biogeochemistry 111(1/3): 661–676.

Marriot CA, Hudson G, Hamilton D, Neilson R, Boag B, Handley LL, Wishart J, Scrimgeour CM, Robinson D (1997) Spatial variability of soil total C and N and their stable isotopes in an upland Scottish grassland. Plant Soil 196: 151–162.

Mooney H, Larigauderie A, Cesario M, Elmquist T, Hoegh-Guldberg O, Lavorel S, Mace GM, Palmer M, Scholes R, Yahara T (2009) Biodiversity, climate change, and ecosystem services. Curr Opin Sust 1: 46–54.

Parker GG (1983) Throughfall and stemflow in the forest nutrient cycle. Adv Ecol Res 13: 57–133.

Pawson SM, Brin A, Brockerhoff EG, Lamb D, Payn TW, Paquette A, Parrota JA (2013) Plantation forests, climate change and biodiversity. Biol Conserv 22: 1203–1227.

Peláez-Silva JA, León-Peláez JD, Lema-Tapias A (2018) Conifer tree plantations for land rehabilitation: An ecological-functional evaluation. Restor Ecol 27: 607–615.

Ramírez JA, León-Peláez JD, Craven D, Herrera DA, Zapata CM, González-Hernández MI, Gallardo-Lancho JF, Osorio W (2014) Effects on nutrient cycling of conifer restoration in a degraded tropical montane forest. Plant Soil 378: 215–226.

Rounsevell MDA, Berry PM, Harrison PA (2006) Future environmental change impacts on rural land use and biodiversity: a synthesis of the ACCELERATES project. Environ Sci Policy 9: 93–100.

Salazar JF, Villegas JC, Rendón AM, Rodríguez E, Hoyos I, Mercado-Bettín D, Poveda G (2018) Scaling properties reveal regulation of river flows in the Amazon through a forest reservoir. Hydrol Earth Syst Sc 22: 1735–1748.

Sardans J, Peñuelas J (2015) Potassium: A neglected nutrient in global change. Global Ecol Biogeogr 24: 261–275.

Scheer MCB (2011) Mineral nutrient fluxes in rainfall and throughfall in a lowland Atlantic rainforest in southern Brazil. J For Res-Jpn 16: 76–81.

Suescún D, Villegas JC, León JD, Flórez CP, García-Leoz V, Correa-Londoño GA (2017) Vegetation cover and rainfall seasonality impact nutrient loss via runoff and erosion in the Colombian Andes. Reg Environ Change 17: 827–839.

Suescún D, León JD, Villegas JC, García-Leoz V, Correa-Londoño GA, Flórez CP (2019) ENSO and rainfall seasonality affect nutrient exchange in tropical mountain forests. Ecohydrology. 2019; e2056.

Tan S, Zhao H, Yang W, Tan B, Ni X, Yue K, Yu Z, Wu F (2018) The effect of canopy exchange on input of base cations in a subalpine spruce plantation during the growth season. Sci Rep-Uk, 8, 9373.

Tropek R, Sedláček O, Keil P, Musilová Z, Símová I, Storch D (2014) Comment on “High resolution global maps of 21st-century forest cover change”. Science 344: 981–981.

Veneklaas EJ (1990) Nutrient fluxes in bulk precipitation and throughfall in two montane tropical rain forests, Colombia. J Ecol 78: 974–992.

Wasige JE, Groen TA, Smaling E, Jetten V (2013) Monitoring basin-scale land cover changes in Kagera Basin of Lake Victoria using ancillary data and remote sensing. Int J Appl Earth Obs 21: 32–42.

